# Advancing Tolerogenic Immunotherapy: A Multi-Epitope Vaccine Design Targeting the CYP2D6 Autoantigen in Autoimmune Hepatitis Through Immuno-Informatics

**DOI:** 10.1101/2024.04.17.589809

**Authors:** Harish Babu Kolla, Anuj Kumar, Roopa Hebbandi Nanjunadappa, Briley Hillyard, Mansi Dutt, Deepak Chauhan, Jean Marshal, David Kelvin, Channakeshava Sokke Umeshappa

## Abstract

Juvenile autoimmune hepatitis (JAIH) is a rare autoimmune disorder affecting children, characterized by the immune system’s misguided attack on liver cells, primarily targeting the CYP2D6 autoantigen. This repeated attack leads to hepatic inflammation, fibrosis, and eventual liver failure. Current therapeutic strategies predominantly rely on immunosuppressive agents or whole B cell depletion antibodies, which render patients susceptible to infections and cancers. Hence, there is an urgent need for antigen-specific therapies to mitigate the severity of autoimmune hepatitis. Tolerogenic antigens represent a promising avenue in immunotherapy, capable of dampening autoimmunity. Here, we present a novel computationally designed multi-epitope tolerogenic vaccine tailored to target CYP2D6, aimed at inducing tolerogenic dendritic cells (DCs) and halting autoimmune progression in JAIH patients. To validate our approach, we have developed a similar vaccine for testing in mouse models of JAIH. The selected tolerogenic epitopes exhibit antigenicity without allergenicity or toxicity, and specifically induce IL-10 production (restricted to CD4+ T cell epitopes). In our vaccine design, tolerogenic poly-epitopes are linked with Toll-like receptor (TLR)-4-agonist, the 50S ribosomal unit, and IL-10, effectively programming DCs towards a tolerogenic state. Molecular docking and dynamic simulations have confirmed strong binding affinities and stable complexes between the vaccine structures, TLR4 and IL-10 receptor alpha (IL-10RA), indicating their potential for *in vivo* DC interaction and programming. Consequently, this innovative vaccine approach demands further exploration through wet lab experiments to assess its tolerogenicity, safety, and efficacy, thereby laying the groundwork for potential application in clinical settings.

## Introduction

Juvenile autoimmune hepatitis (JAIH), also known as type 2 AIH, poses a significant threat to children. It is a life-threatening inflammatory condition characterized by interface hepatitis and the gradual deterioration of liver tissue due to the immune system’s attack, culminating in chronic fibrosis, liver failure, and potentially, mortality [1-3]. JAIH predominantly affects girls from infancy through late adolescence[4]. Globally, the World Health Organization notes a concerning rise in the incidence and prevalence of AIH compared to other autoimmune liver diseases (AILDs) [5]. In the USA alone, a projected 100,000 to 200,000 persons are affected by AIH. Numerous factors influence the onset and progression of AIH in children, with high-risk genetic elements playing a pivotal role. Genome-wide association studies (GWAS) have unequivocally linked specific HLA alleles, notably HLA-DRB1*03:01 and HLA-DRB1*07:01, with type 2 AIH [2, 6]. This genetic association underscores the crucial involvement of antigen-presenting cells (APCs) and activated CD4+ T helper cells in the pathogenesis of the disease.

In a healthy state, Antigen Presenting Cells (APCs) typically present pathogenic epitopes, from foreign antigens, to CD4+ and CD8+ T immune cells. However, in the case of JAIH, APCs process autoantigens like cytochrome p450 2D6 (CYP2D6) and present these epitopes to autoreactive T cells. Consequently, these T cells launch an attack on liver tissue, contributing to liver failure [7]. Proper and effective treatment of JAIH is essential for controlling the severity of AIH. Presently, immunosuppressive drugs are utilized to suppress the immune response in the body [8]. However, these drugs primarily alleviate symptoms and do not effectively address the underlying disease. Another treatment approach for autoimmune diseases involves depleting total B cells, including autoreactive B cells, using anti-CD20 monoclonal antibodies [9]. While this method eliminates autoreactive B cells, it also affects normal B cells crucial for overall immune function [10]. To address the limitations of nonspecific immune modulating therapies, the development of antigen-specific tolerogenic immunotherapies is imperative. These therapies have the potential to halt disease progression without compromising systemic immunity [11][12-15].

Recent advancements in computational technologies have revolutionized the field of immunotherapy. These tools enable epitope mapping, structural design, and molecular interaction studies critical for effective vaccine development. Leveraging immunoinformatic approaches, numerous vaccines have been successfully developed against various infections and cancers [16-21]. However, the application of such technologies in treating autoimmune diseases remains relatively unexplored. In this study, we harnessed these advanced technologies to predict autoreactive CD4+ T cell epitopes within the CYP2D6 autoantigen and successfully designed a multi-epitope-based tolerogenic vaccine using these predicted epitopes. The tolerogenic vaccine incorporates an adjuvant to activate APCs, such as dendritic cells (DCs), and a tolerogenic cytokine, IL-10, capable of converting the phenotype of activated APCs into tolerogenic ones. To assess the structural stability of the designed vaccine models, we conducted Ramachandran plot analysis. Furthermore, we evaluated their potential to interact with toll-like receptors (TLRs) on APCs and IL-10 receptor alpha (IL-10RA) through *in silico* molecular docking and dynamic simulation studies. Therefore, further wet lab experiments are necessary to evaluate the vaccine’s tolerogenicity, safety, and efficacy, thereby setting the stage for potential clinical application.

## Methodology

### Sequence retrieval

The complete amino acid sequence of the mouse and human CYP2D6 autoantigen with the corresponding accession IDs (Q91W87; P10635) were retrieved from Uniprot resource. These amino acid sequences were used for the *in silico* prediction of CD4+ T cell epitopes and further downstream analysis.

### Epitope prediction

The CD4+ T cell epitopes were predicted in amino acid sequences of mCYP2D6 and hCYP2D6 autoantigens. The epitopes were predicted using NetMHCIIpan 4.1 BA module in the IEDB Analytical resource (http://tools.iedb.org/mhcii/) [22]. While predicting the epitopes, H2-IAd allele was selected for mouse and the high-risk HLA alleles-HLA-DRB1^*^03:01 and HLA-DRB1^*^07:01 were selected for human. The epitopes were predicted based on the binding affinity between the the allele/MHC II molecule and the peptide. These epitopes were screened for the presence of non-allergic, non-toxic, non IFNLJ inducers and IL-10 inducers for the selection of tolerogenic epitopes.

### Screening of tolerogenic epitopes

For the identification of tolerogenic epitopes, firstly we subjected the predicted epitopes for their allergenicity profile. Allergenicity of these epitopes was predicted using AllerTop v 2.0 server (https://www.ddg-pharmfac.net/AllerTOP/) [23]. The predicted non-allergenic epitopes were further screened for the non-toxic epitopes by using ToxiPred web server (https://webs.iiitd.edu.in/raghava/toxinpred/) [24]. Once we identified non-allergic and non-toxic epitopes, the tolerogenic epitopes were finalized based on their ability to induce IL-10 and being non-IFNLJ inducers. The IFN□inducing epitopes were identified by submitting the epitope peptide sequences to IFNepitope server (https://webs.iiitd.edu.in/raghava/ifnepitope/predict.php) [22] and and IL-10 inducing epitopes with IL-10Pred sever (https://webs.iiitd.edu.in/raghava/il10pred/algo.php) respectively [25].

### Design of tolerogenic vaccine candidates

We have designed two vaccine candidates targeting mouse and human CYP2D6 autoantigen. The screened tolerogenic epitopes were included in the final vaccine design. We have included a 50 S ribosomal subunit as an adjuvant at the N-terminal end of the vaccine to activate dendritic cells. The adjuvant was separated from the vaccine with an EAAAK linker. On the other hand, IL-10 was also included in the vaccine construct to induce tolerance against the CYP2D6 autoantigen. The IL-10 cytokine was separated from the epitopes by a flexible GGGGS linker. The CD4+ tolerogenic T cell epitopes were separated by AAY linkers as reported by previous studies for the design of a multi-epitope peptide vaccine. After the vaccine design, the vaccine constructs were evaluated for their physicochemical properties using ProtParam tool (https://web.expasy.org/protparam/).

### Protein modeling and refinement

Three dimensional tertiary structures of the mCYPD-tol vac and hCYPD-tol vac were predicted *in silico* for molecular docking studies. The 3-D structure of mCYPD-Tol vac and hCYPD-Tol vac proteins were predicted based on their amino acid sequence and this step is called protein modeling. Protein modeling was carried out using Robetta server (https://robetta.bakerlab.org/). The predicted structures were further refined using GalaxyRefine web server [26] (https://galaxy.seoklab.org/cgi-bin/submit.cgi?type=REFINE).

### Molecular docking

Molecular docking studies were carried out to determine the interactions between the vaccine components and their corresponding responses during innate immune response. Firstly, we performed molecular docking between the vaccine candidates and TLR4. This enables us to understand the interaction of 50 S ribosomal subunit adjuvant protein with TLR4 as a part of dendritic cell activation. Later, the vaccine constructs were docked with the IL-10RA to determine whether the IL-10 cytokine component included in the vaccine candidates can successfully interact with the IL-10RA, which is very crucial in the conversion of the phenotype of autoantigen specific dendritic cells into tolerogenic dendritic cells (DCs).

While performing the molecular docking studies, the target receptors (TLR4 and IL-10RA) were designated as chain A and the vaccine construct ligands as chain B. Later, the two-dimensional amino acid interactions in terms of total number of H-binds, salt bridges and non-bonded interactions were visualized in the docked complexes through PDBsum sever (https://www.ebi.ac.uk/thornton-srv/databases/pdbsum/) [27] and the binding affinity between the ligand and receptor in a docked complex that are mediated by these interactions was predicted using Prodigy Web server (https://wenmr.science.uu.nl/prodigy/) [28].

### Molecular dynamics simulations

MD simulations were carried out for all the docking complexes (mCYPD-Tol Vac-TLR4, mCYPD-Tol vac-IL-10RA, hCYPD-Tol Vac-TLR4 and hCYPD-Tol vac-IL-10RA) to determine the stability of the interactions at atomic level. We performed MD simulations for 100 nanoseconds similar to previous studies [29-31]. The AMBER99SB-ILDN protein and nucleic AMBER94 force fields of GROMACS 2023 package were employed to carry out the MD simulations. All of the four docked complexes were solved using the transferable intermolecular potential 3P (TIP3P) water model and equilibrated with NVT ensemble at 300K and the Parinello-Rahman barostat coupling ensembles.

## Results

### Prediction and screening of epitopes

The CD4+ T cell epitopes in mouse and human CYP2D6 autoantigens were predicted based on the binding affinity and IC50 value (nM) between the epitope and the corresponding MHC II or HLA II alleles. Only epitopes with an IC50 value less than or equal to 1000 nM have been considered. Considering this criterion, we obtained a total of 9 and 190 CD4+ T cell epitopes in mouse and human CYP2D6 autoantigens, respectively **[Figure 1A, B; Supplementary table (ST) 1 and 2]**. These predicted epitopes were further screened based on the allergenicity, toxicity, and ability to induce IL-10 cytokines. After subjecting these predicted epitopes for a robust screening, we obtained 2 epitopes corresponding to mouse and 82 epitopes for human systems to include in the design of a tolerogenic vaccine **(Figure 1A, B; Table S1)**.

**Figure 1.**
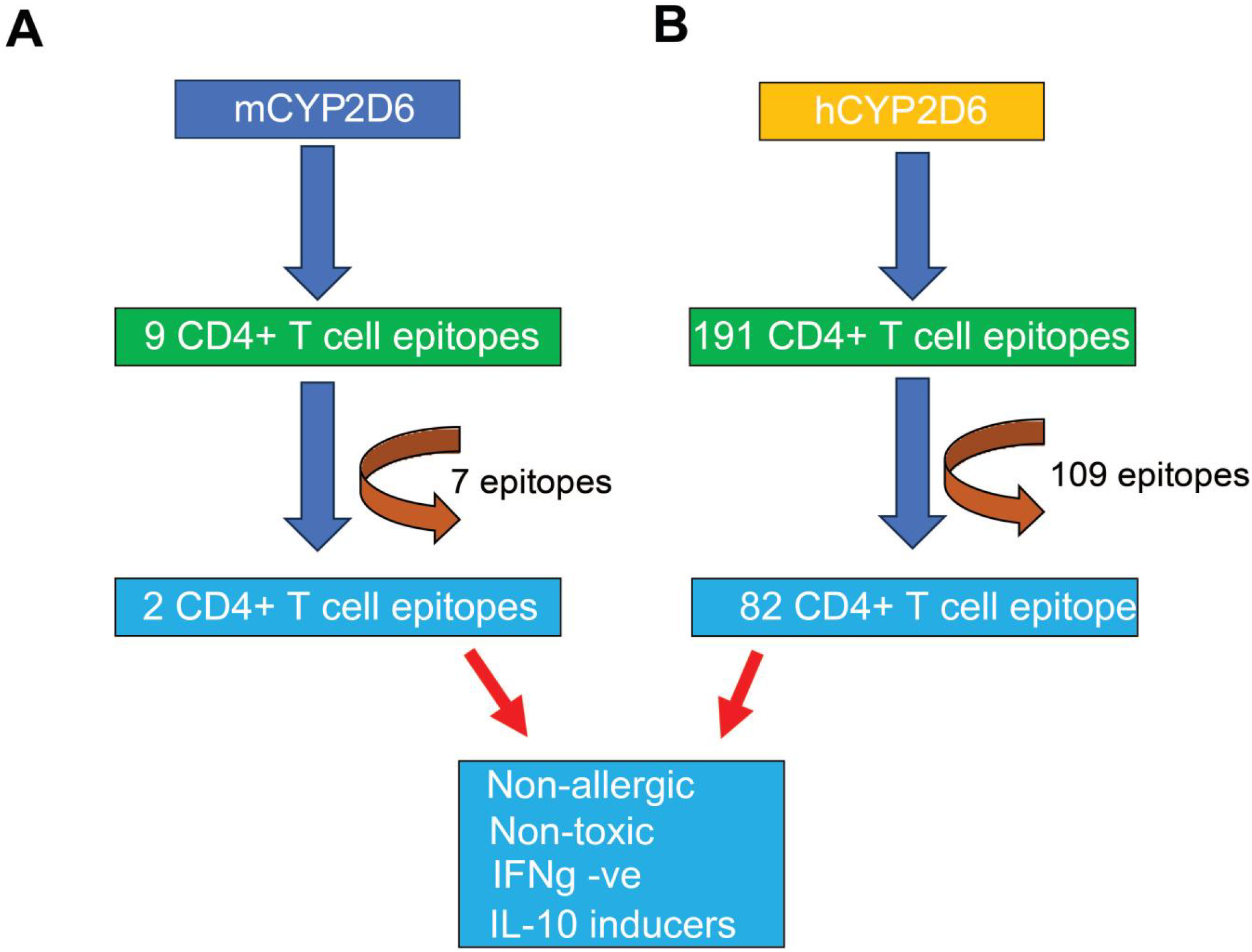
Overview of the design of tolerogenic multi-epitope vaccine targeting CYP2D6 autoantigen. **A, B**. Flow chart representing the screening of CD4+ T cell epitopes for the design of tolerogenic vaccine targeting the mouse CYP2D6 (A) and human CYP2D6 (B).

### Design of tolerogenic vaccines

We have designed two tolerogenic vaccines (one for mouse to obtain a proof-of-concept in laboratory studies and one for human) based on the obtained epitopes after a thorough screening. Interestingly we found that most of the epitopes are overlapping or adjacent. Thus, we designed the vaccines including those immunodominant regions or overlapping epitopes in both mouse and human systems. For example, we identified 2 epitopes in mice overlapping between 302-319 amino acids in mCYPD2D6 autoantigen **(Figure 2A)**. Similarly, in the case of human CYP2D6, the tolerogenic epitopes corresponding to both the considered HLA alleles were present in the regions 189-255, 292-310 and 386-412 amino acids **(Figure 2B)**. These sequences were finally considered for the design of a tolerogenic vaccine for JAIH.

**Figure 2.**
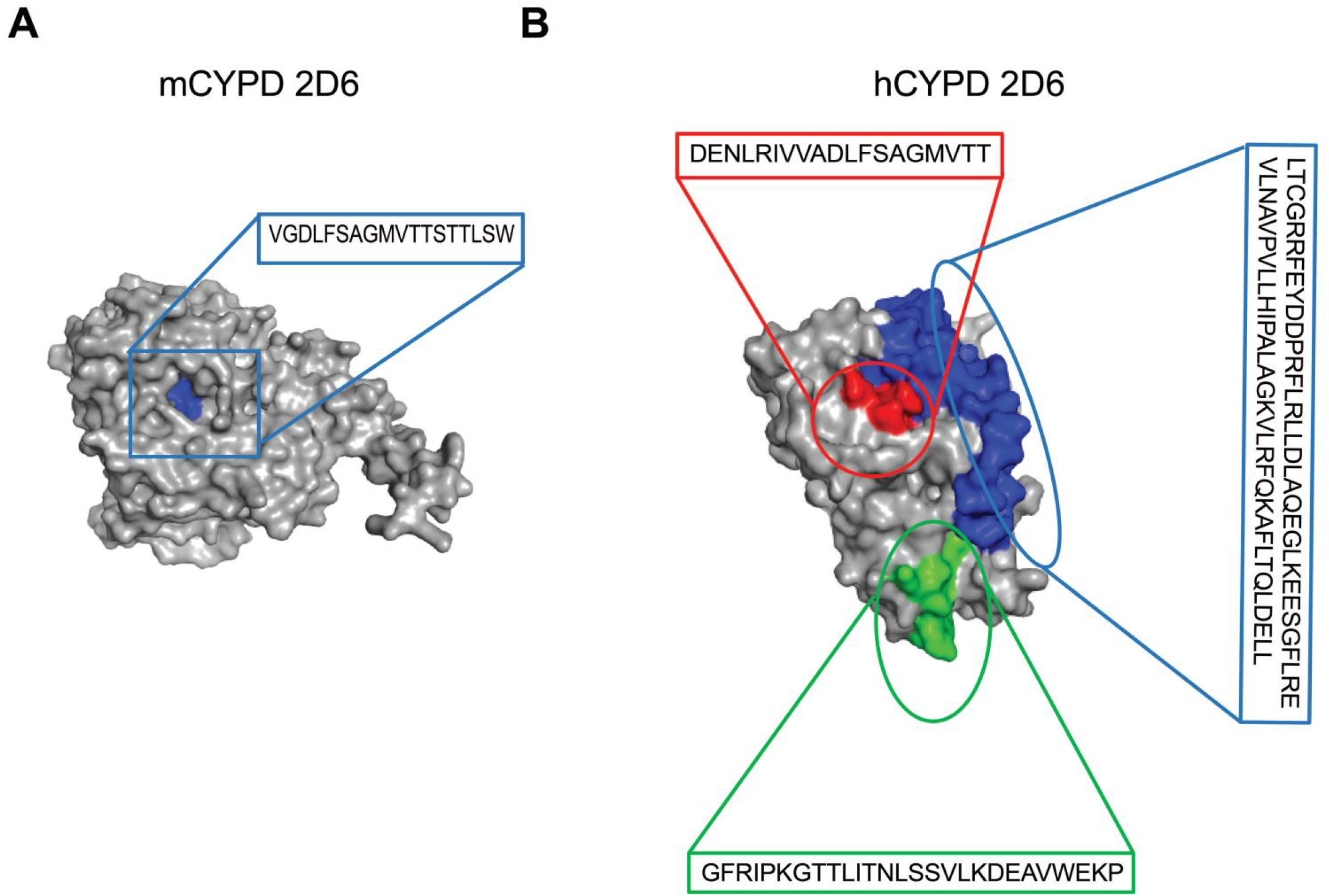
Tolerogenic multi-epitope regions selected for vaccine design in the CYPD2D6 autoantigens. **A, B**. Representation of tolerogenic multi-epitope region in the mouse CYP2D6 with overlapping CD4+ T cell epitope (highlighted in blue) (A) and human CYP2D6 autoantigen with overlapping CD4+ T cell epitopes (highlighted in red, blue and green colours) (B) for the design of tolerogenic multi-epitope vaccine. Note: *In silico* protein structure predictions were generated using the Robetta server.

During vaccine design, a 50S ribosomal subunit adjuvant was incorporated at the N-terminal end of the construct to facilitate interaction with and activation of dendritic cells (DCs) via TLR4 signaling. This adjuvant was linked to the vaccine peptides by an EAAAK linker sequence **(Figure 3A, B)**. Similarly, the immunogenic peptides in the vaccine construct were separated by AAY linker, which aid in processing and presentation of epitope peptides to the CD4+ T helper cells. On the other hand, an anti-inflammatory cytokine IL-10 was added at the C-terminal end of the vaccine construct to induce tolerance against CYP2D6 autoantigen **(Figure 3A, B)**. The IL-10 was separated from the vaccine construct with the help of a flexible GGGGS linker. The designed vaccine constructs were then used for further analysis.

**Figure 3.**
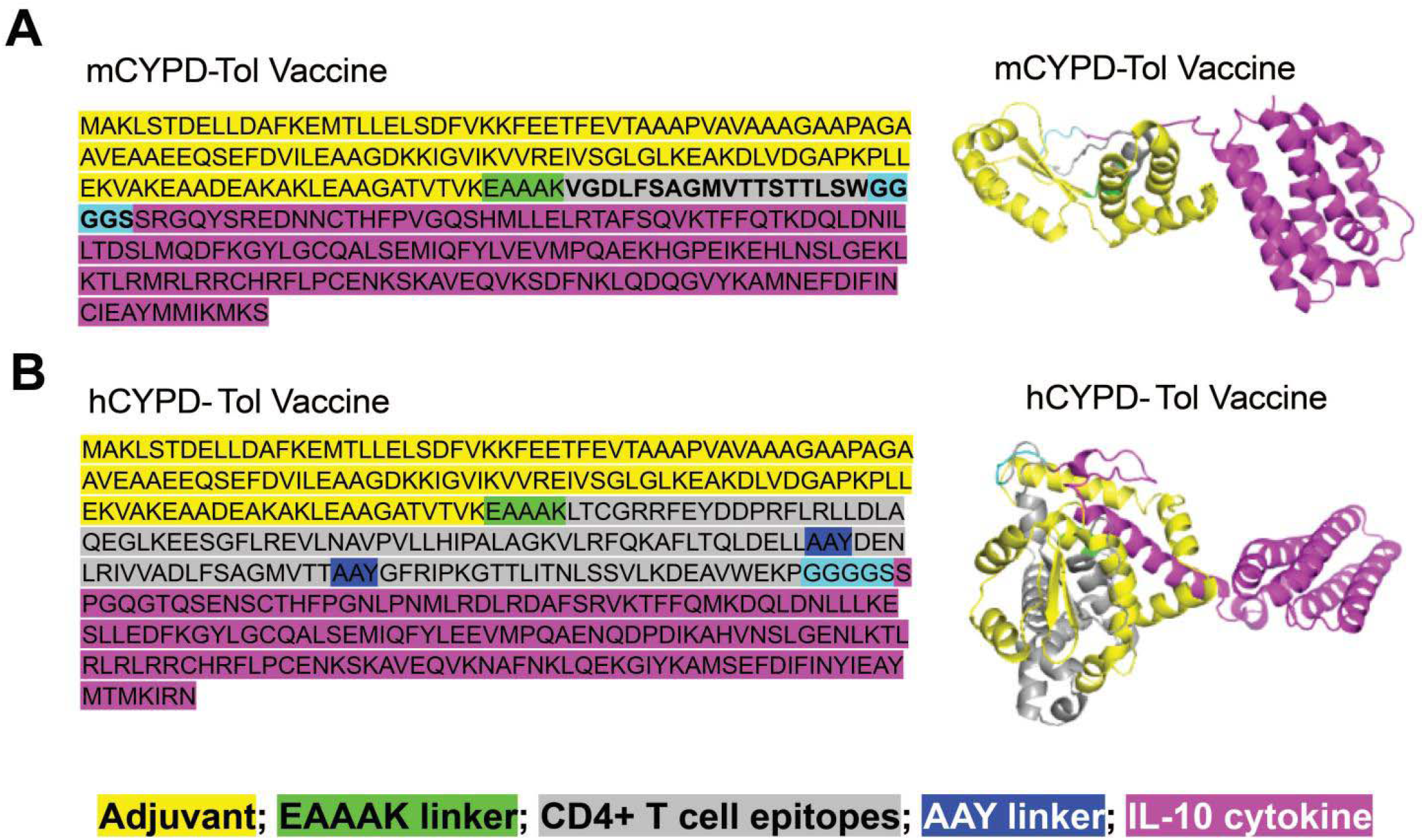
Sequence and structure of multi-epitope vaccines. **A, B**. Design of mouse tolerogenic CYP2D6-relevant multi-epitope vaccine specific mouse (A) and human (B), showing amino acid sequences (left) and coiled-coil protein structures (right) generated through protein modeling with Robetta server. The 50S ribosomal subunit (yellow) is used as an adjuvant for the vaccine construct (grey) Whereas the IL-10 cytokine is fused to the C-terminal domain (pink) of the vaccine construct to induce tolerance.

### Evaluation of the vaccine constructs

The designed vaccine constructs were evaluated for their physiochemical attributes. We evaluated the physicochemical properties of tolerogenic vaccines, such as molecular weight, theoretical pI, instability index, aliphatic index, and Grand average of hydropathicity (GRAVY). The computed values of these parameters for the mouse tolerogenic vaccine are 34817.07, 5.37, 35.18, 85.66 and -0.156, respectively. Similarly, these values corresponding to the human tolerogenic vaccine are 46161.19, 5.26, 37.83, 94.84 and -0.128, respectively. Both the vaccines were stable based on the instability index values, which is below 40 for stable proteins.

### Protein modeling and evaluation

Three dimensional structures were predicted for the designed tolerogenic vaccine candidates for molecular docking studies. The predicted structures were further refined to improve their RMSD values. The structural stability of the protein structures of vaccine constructs was evaluated by a popular method called Ramachandran plot analysis. As a general criterion, any protein model whose Ramachandran plot comprises ∼90 % of the total amino acid residues falling in most favoured regions are considered to be of good quality for structural studies or molecular docking. Similarly, the mouse and human tolerogenic vaccine constructs comprise of 92.3 % and 91.2 % of total amino acid residues in favourable region, 7.3 % and 7.4 % in additionally allowed regions with only few remaining residues in generously allowed and disallowed regions **(Figure 4 A, B)**.

**Figure 4.**
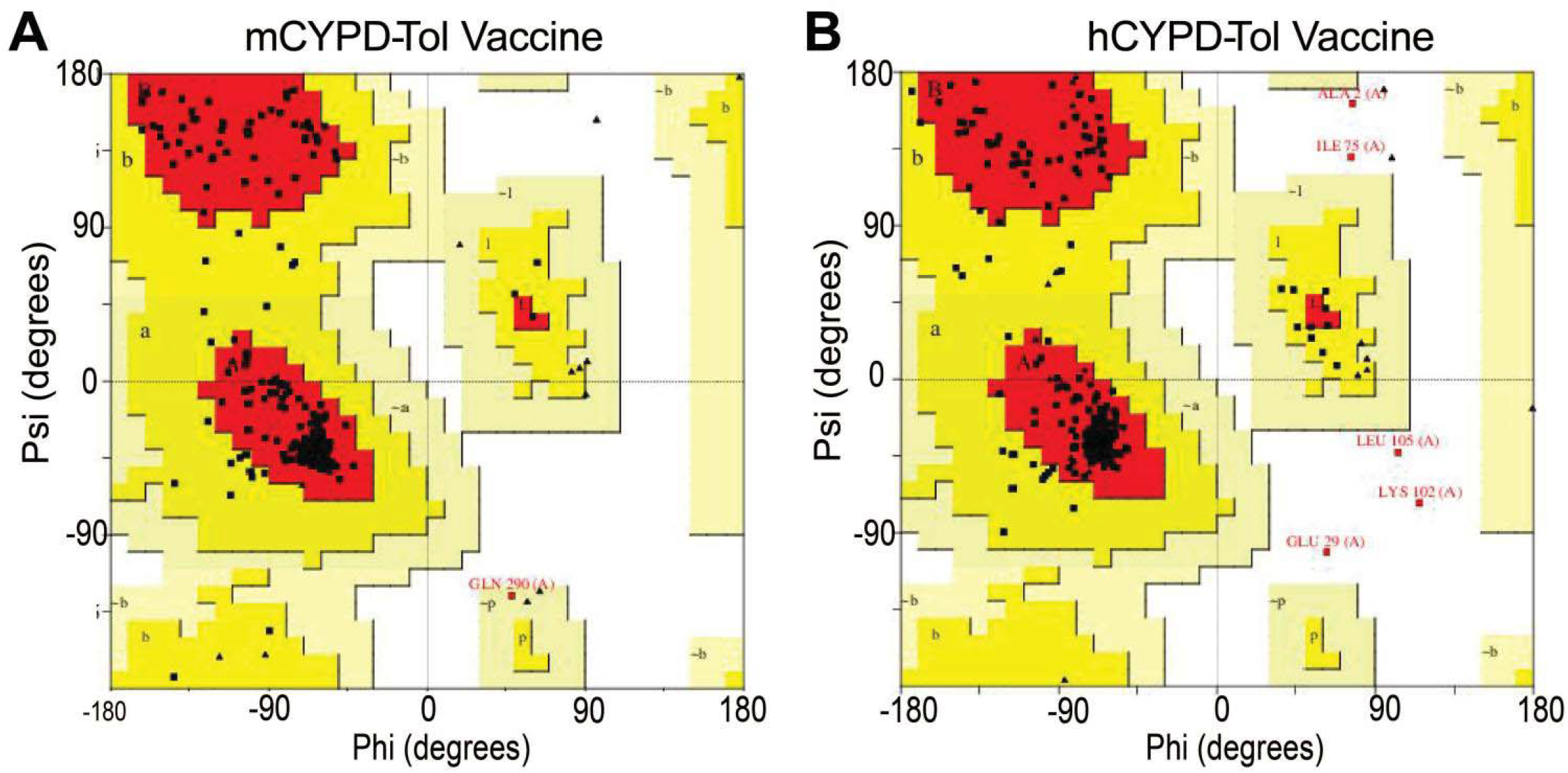
Evaluation of three-dimensional strcutures of the tolerogenic vaccine constructs. **A, B**. Ramachandran plot diagrams showing the overall structural quality of mouse CYP2D6 (A) and human CYP2D6 (B) tolerogenic vaccines.

### Molecular docking

Molecular docking is a widely used computational method to study the interaction between two biomolecules. It is being considered to study the interaction between vaccine and target protein receptors. Likewise, we studied the interaction between the components in the designed tolerogenic vaccines and the target receptors through molecular docking studies which enabled us to understand the potential role of adjuvant and IL-10 components in inducing the immune tolerance against the CYP2D6 autoantigen. The first interaction that we studied is between the TLR4 and the 50 S ribosomal subunit adjuvant in mouse tolerogenic vaccine where we observed a strong affinity between the two molecules with a Gibbs free energy (G) value of -11.6 KCal/mol which is stabilized by 6 H-bonds, 3-Salt bridges and 272 non-bonded interactions **(Figure 5A)**. Similarly, both the TLR4 and adjuvant in human tolerogenic vaccine strongly interacted through 5 H-bonds, 4 salt bridges and 242 non-bonded interactions with a binding affinity of -12 KCal/mol **(Figure 5B)**.

**Figure 5.**
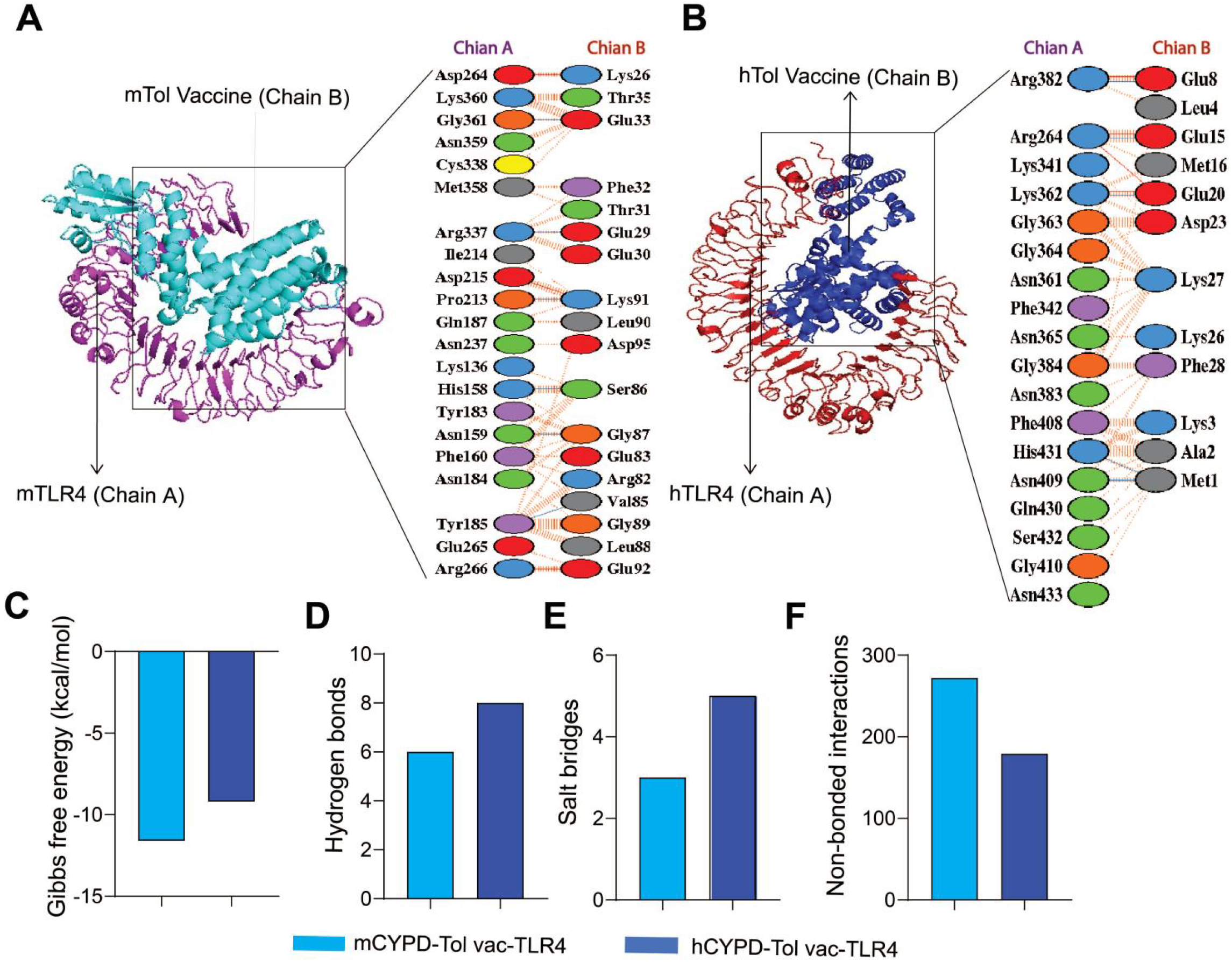
Molecular docking and interactions between the 50S ribosomal subunit with mouse and human TLR4 structures. **A**. Visualization of 2D and 3D molecular interactions between the 50S ribosomal subunit in the mouse tolerogenic vaccine (Chain B) and TLR4 (Chain A). **B**. Visualization of the 2D and 3D molecular interactions between the 50S ribosomal subunit in the human tolerogenic vaccine (Chain B) and TLR4 (Chain A). **C**. Gibbs free energy. **D**. Hydrogen bonds. **E**. Salt bridges. **F**. Non-bonded interactions. The Bar graph representing the overview of 2-D interactions between the two molecules in the docked complex, showing Gibbs free energy (C), H-bonds (D), Salt bridges (E), and non-bonded interactions (F).

Secondly, we also explored the interactions between the IL-10 component included in the vaccine design and the IL-10RA. The IL-10 in the mouse tolerogenic vaccine bound very strongly with the IL-10 receptor forming 8 H-bonds, 5 salt bridges and 179 non bonded interactions contributing to a G value of -9.2 KCal/mol **(Figure 6A)**. Additionally, the IL-10 component within the human tolerogenic vaccine exhibited a comparable binding affinity of -9.5 kcal/mol with the IL-10RA. This interaction was facilitated by 5 hydrogen bonds, 2 salt bridges, and 265 non-bonded interactions **(Figure 6B)**.

**Figure 6.**
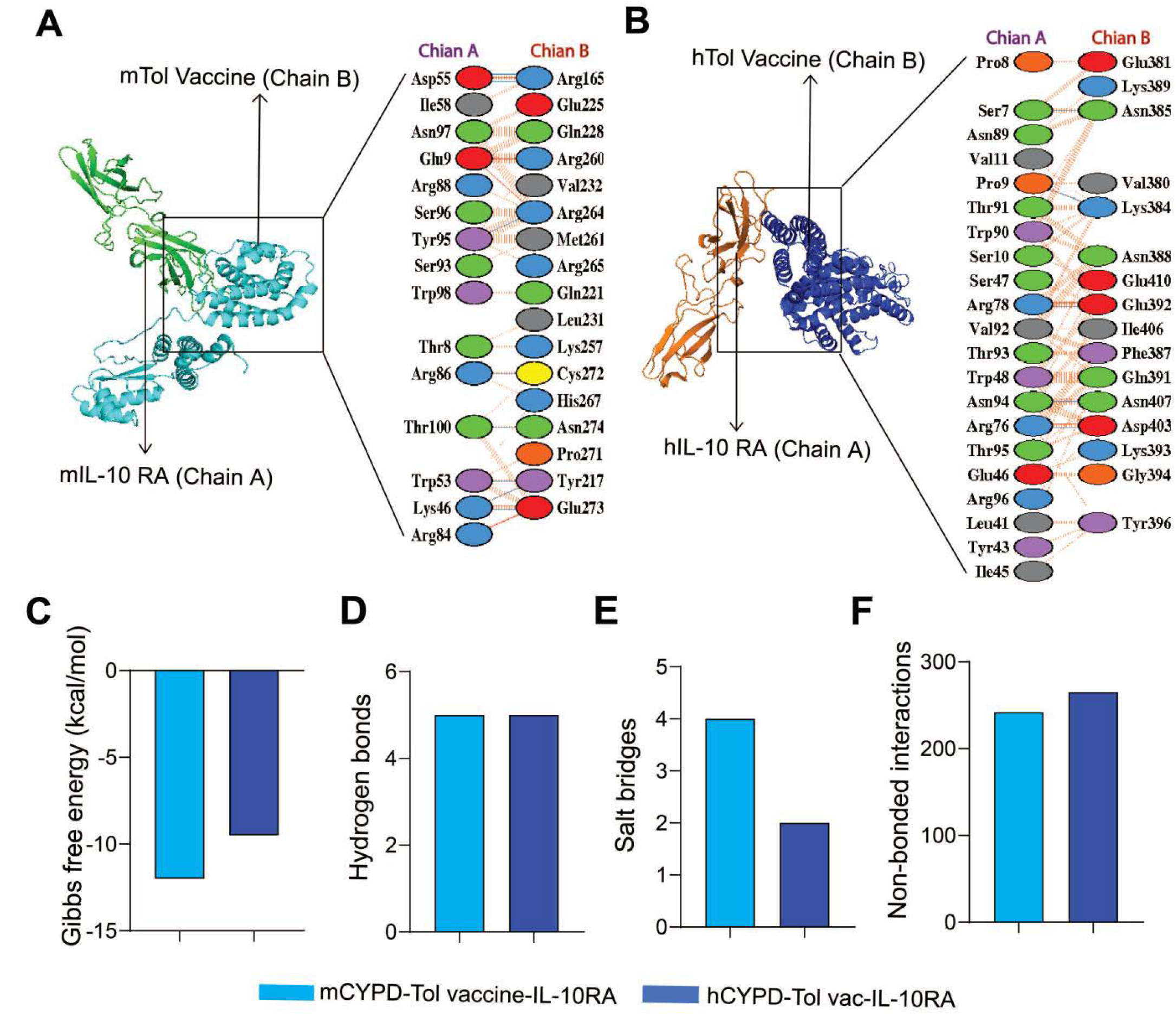
Molecular docking and interactions between the IL-10 cytokine with mouse and human IL-10RA structures. **A**. Visualization of the 2D and 3D molecular interactions between the IL-10 cytokine in the mouse tolerogenic vaccine (Chain B) and IL-10RA (Chain A) and **B**. Visualization of the 2D and 3D molecular interactions between the IL-10 cytokine in the human tolerogenic vaccine (Chain B) and IL-10RA (Chain A). **C**. Gibbs free energy. **D**. Hydrogen bonds. **E**. Salt bridges. **F**. Non-bonded interactions. The Bar graph representing the overview of 2-D interactions between the two molecules in the docked complex, showing Gibbs free energy (C), H-bonds (D), Salt bridges (E), and non-bonded interactions (F).

### MD simulations

The dynamic behavior of human and mouse vaccines docked complexes with 10RA and TLR4 receptors were assessed using calculating the root mean square deviation (RMSD) and root mean square fluctuation (RMSF) plots using the GROMACS package on 100 ns.

RMSD plot analysis is one of the popular approaches to calculating the changes in the simulated systems during MD simulations. To investigate the stability of the docked complexes at the atomic level, RMSDs of the vaccine and receptors docking complexes were calculated and graphically investigated. The average RMSD values for the hCYPD-Tol vac-IL-10RA (blue), hCYPD-Tol vac-TLR4 (red), mCYPD-Tol vac-IL-10RA (green), and mCYPD-Tol vac-TLR4 (purple) docked complexes were 1.09, 0.72, 0.83, and 0.60 nm, respectively. The docking complexes of vaccines and receptors exhibited the stability pattern throughout the simulation on 100 ns with a range between ∼0.40 and ∼1.70 nm. As evident from **Figure 7A**, the complex of hCYPD-Tol vac-IL-10RA (blue) showed two major fluctuations between ∼21 to ∼43 ns and ∼44 to ∼63 ns. After 63 ns, this complex demonstrated stability up to 100 ns on ∼1.0 nm. The other three complexes, namely, hCYPD-Tol vac-TLR4 (red), mCYPD-Tol vac-IL-10RA (green), and mCYPD-Tol vac-TLR4 (purple) showed the almost same pattern of stability on a 100 ns time scale. No major fluctuations have been noted in these complexes. It can be concluded from the calculated RMSDs that docking complexes showed minimal conformational changes and are stable at the atomic level.

**Figure 7.**
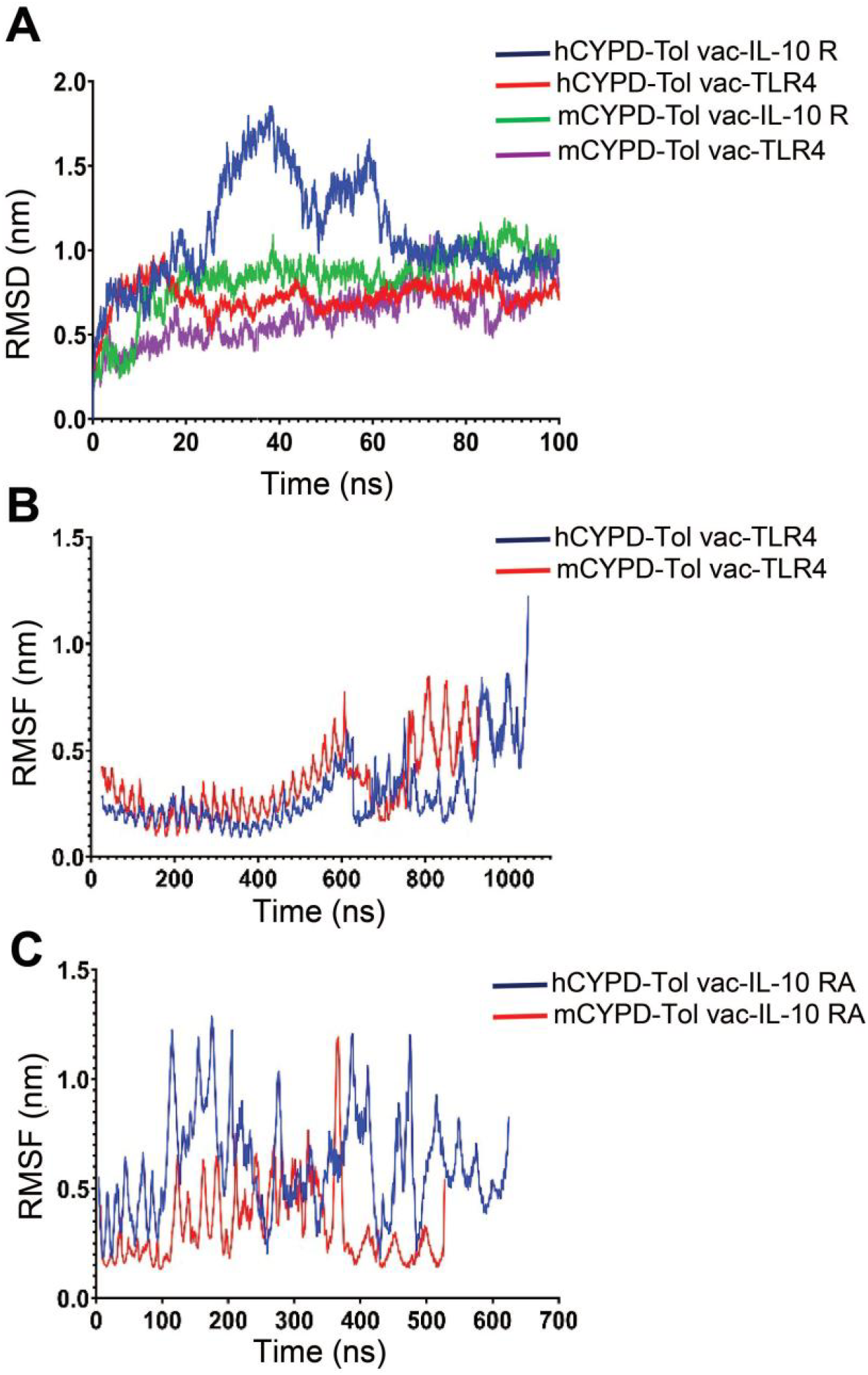
Molecular dynamics simulations representing the stability of the Docking Complexes. **A**. Line diagram displaying the calculated RMSD values in the docking complexes between the mouse and human vaccine components (50S ribosomal subunit or IL-10) and corresponding host receptors (TLR-4 or IL-10RA, respectively. **B**. Line diagram illustrating the RMSF values of the docking complexes between mouse and human vaccine component, 50S ribosomal subunit adjuvant, and corresponding species-specific TLR-4. **C**. Line diagram illustrating the RMSF values of the docking complexes between mouse and human vaccine component, IL-10, and corresponding species-specific IL-10RA.

To calculate the individual residue flexibility of simulated systems, RMSF plot analysis was performed for all four docking complexes. The average RMSF values for the hCYPD-Tol vac-IL-10RA (blue), hCYPD-Tol vac-TLR4 (red), mCYPD-Tol vac-IL-10RA (green), and mCYPD-Tol vac-TLR4 (purple) docked complexes were 0.60, 0.29, 0.32, and 0.33 nm, respectively. As shown in **Figure 7B**, the complex of hCYPD-Tol vac-IL-10RA (blue) showed several fluctuations throughout the simulations on a 100 ns time scale. The major fluctuations have been noted between 100 to 200 residues, and 390 to 490 residues **(Figure 7C)**. The complex of mCYPD-Tol vac-IL-10RA (red) exhibited the highest peak between 359 to 363 residues. The Human and Mouse vaccine complexes with TLR4 receptor molecule demonstrated an almost similar pattern of peaks throughout the simulations. In these complexes, the highest peak was observed at the end in the hCYPD-Tol vac-TLR4 complex (blue). Calculated RMSF plots indicated that all four designed vaccine constructs significantly interact with their corresponding receptor molecules.

## Discussion

In this study, we developed and computationally assessed a multi-epitope tolerogenic vaccine directed at the CYP2D6 autoantigen implicated in JAIH development. Unlike traditional immunogenic vaccines that are designed to prevent infections, these tolerogenic vaccines aim to dampen immune responses [32-35]. During the vaccine design, careful screening of potential CD4+ T cell tolerogenic epitopes was conducted, considering allergenicity, toxicity, and IL-10-inducing properties. Structural modeling of the vaccine constructs demonstrated stability for molecular docking studies. Molecular docking and MD simulations confirmed the potential of both vaccine constructs to induce immune tolerance. Strong and stable interactions between the TLR4 and 50S ribosomal subunit adjuvant, mediated by numerous H bonds, salt bridges, and non-bonded interactions, which suggest potential activation of DCs *in vivo* by the vaccine constructs. Similarly, observed interactions between IL-10 and IL-10RA indicate IL-10’s ability to provide tolerogenic signals to DCs. MD simulations further validated the stability of interactions within the docked complexes, supporting potential interactions of vaccine candidates with DCs *in vivo*.

Our previous work has demonstrated the efficacy of antigen-specific tolerogenic immunotherapies employing peptide-MHC-II-based nanomedicines [12-15]. These therapeutics, containing tissue-specific autoantigenic epitopes, have successfully reprogrammed autoantigen-experienced activated CD4+ T-cells into IL-10-secreting T-regulatory type-1 cells (TR1). This transformation fosters a local immunosuppressive environment and facilitates the conversion of local dendritic cells (DCs) into tolerogenic DCs, thereby impeding the progression of autoimmune pathologies. Importantly, these regulatory T cells do not compromise local or systemic immunity against infection and cancer, indicating their safety for use in autoimmune patients [12, 14, 15]. The underlying reprogramming mechanisms are largely orchestrated by the regulatory cytokine IL-10. Specifically, interactions between regulatory T cells and APCs stimulate local IL-10 production, which suppresses antigen presentation, T cell activation, and recruitment [15]. Moreover, IL-10 production via TR1 interactions promotes B-cell differentiation into tolerogenic APCs [12, 14], effectively halting the generation of autoreactive T and B cells and thereby controlling autoimmunity.

In contrast, the vaccine developed in this study directly targets dendritic cells (DCs), aiming to activate and convert their phenotype into tolerogenic DCs. The inclusion of the 50S ribosomal unit adjuvant serves to activate DCs, facilitating the uptake, processing, and presentation of autoantigenic epitopes to CD4+ T cells. Specifically, the 50S ribosomal protein L7/L12 derived from Mycobacterium tuberculosis has been shown to induce DC maturation and T cell activation [36]. This adjuvant is well-recognized by Toll-like receptor 4 (TLR4) during DC maturation. However, in the presence of the regulatory cytokine, IL-10, incorporated into the vaccine, DCs undergo reprogramming to adopt a tolerogenic phenotype [37-39]. Consequently, activated CD4+ T cells differentiate into tolerogenic regulatory CD4+ T cells (TR1 cells), establishing a local regulatory network in the liver. IL-10 is an anti-inflammatory cytokine crucial for inducing tolerance [40]. Collectively, the incorporation of these components in the vaccine construct is expected to effectively induce antigen-specific immune tolerance against the CYP2D6 autoantigen.

Previous research has underscored the therapeutic potential of antigen-specific tolerogenic immunotherapies in various autoimmune diseases, including rheumatoid arthritis, multiple sclerosis, and type-1 diabetes [41, 42]. Tolerogenic DCs have also shown promise in promoting liver transplant tolerance [43]. However, existing studies have primarily focused on in vitro-generated tolerogenic DCs, which are labor-intensive and costly. Our vaccine, capable of generating tolerogenic DCs *in vivo*, offers a promising alternative for treating autoimmune diseases, including JAIH, in an antigen-specific manner—a longstanding goal in autoimmune therapy.

In conclusion, our study presents a novel application of tolerogenic DCs through the development of a tolerogenic polypeptide vaccine capable of converting endogenous DCs into tolerogenic ones in an antigen-specific manner. These findings not only highlight the potential of our vaccine in inducing antigen-specific immune tolerance against the CYP2D6 autoantigen in JAIH but also validate the use of in-silico techniques for designing such vaccines against other autoimmune conditions. Future research endeavors should prioritize experimental investigations aimed at evaluating the immune suppressive nature and mechanisms underlying antigen tolerance induced by our vaccine constructs. As current JAIH treatment predominantly relies on systemic immunosuppressive therapies, the development of disease-specific tolerogenic vaccines offers a promising avenue for non-immunosuppressive treatment options that spare local and systemic immunity intact against infections and cancers. As research progresses, our novel in-silico model will continue to play a pivotal role in providing a flexible approach for designing tolerogenic multi-epitope peptide vaccines, not only for JAIH but also for other autoimmune diseases.

## Author contributions

H.B.K generated data in figures 1-6 with the contributions from A.K, B.H, R.H.N and D.C under the supervision of C.S.U. A.K and M.D. generated figure 7 under the supervision of D.K. and C.S.U. B.H, R.H.N and D.C contributed to data interpretation, figure preparations (Fig 1, 2, 3 or 4), and some sections of the manuscript writing under the supervision of C.S.U. and/or J.M. H.B.K wrote the first draft with the help of C.S.U and A.K. C.S.U. designed the study, supervised, and coordinated its execution, and wrote the manuscript along with H.B.K and A.K.

## Acknowledgments

H.B.K is supported by graduate studentship by DMRF. R.H.N is supported by the fellowships from the Immunity, Infection, Inflammation and Vaccinology (I3V) and American Association of Immunologists organizations. D.C is supported by the postdoctoral fellowships from IWK and I3V. This work is supported by Dalhousie University start-up funding and prestigious Canada Research Chair Tier 2 Award from Canadian Institutes of Health Research.

