## Supplementary tables for "Advancing Tolerogenic Immunotherapy: A Multi-Epitope Vaccine Design Targeting the CYP2D6 Autoantigen in Autoimmune Hepatitis Through Immuno-Informatics"

**Supplementary table 1.** MHC II specific CD4+ T cell epitopes predicted in the CYP 2D6 autoantigen.

| **Allele** | **Peptide Sequence** | **IC50** |
| --- | --- | --- |
| H2-IAb | GDLFSAGMVTTSTTL | 214.67 |
| H2-IAb | VGDLFSAGMVTTSTT | 227.03 |
| H2-IAb | VVGDLFSAGMVTTST | 255.90 |
| H2-IAb | DLFSAGMVTTSTTLS | 296.22 |
| H2-IAb | LFSAGMVTTSTTLSW | 571.86 |
| H2-IAb | QGHFVKPEAFMPFSA | 841.99 |
| H2-IAb | GHFVKPEAFMPFSAG | 885.99 |
| H2-IAb | VQRFADILPLGVPHK | 958.45 |
| H2-IAb | HFVKPEAFMPFSAGR | 992.51 |

**Supplementary table 2.** HLA II specific CD4+ T cell epitopes predicted in CYP 2D6 autoantigen.

| **Allele** | **Peptide Sequence** | **IC50** |
| --- | --- | --- |
| HLA-DRB1*07:01 | AACLCAAFANHSGRP | 587.83 |
| HLA-DRB1*07:01 | AAFANHSGRPFRPNG | 247.36 |
| HLA-DRB1*07:01 | ACLCAAFANHSGRPF | 206.74 |
| HLA-DRB1*07:01 | AFANHSGRPFRPNGL | 417.67 |
| HLA-DRB1*07:01 | AFLPFSAGRRACLGE | 237.19 |
| HLA-DRB1*07:01 | AFLVSPSPYELCAVP | 242.63 |
| HLA-DRB1*07:01 | AGMVTTSTTLAWGLL | 176.51 |
| HLA-DRB1*03:01 | AIFLLLVDLMHRRQR | 30.40 |
| HLA-DRB1*07:01 |  | 713.47 |
| HLA-DRB1*03:01 | AQEGLKEESGFLREV | 530.33 |
| HLA-DRB1*07:01 | AQGHFVKPEAFLPFS | 429.30 |
| HLA-DRB1*07:01 | ARMELFLFFTSLLQH | 459.23 |
| HLA-DRB1*07:01 | AVPVLLHIPALAGKV | 57.96 |
| HLA-DRB1*07:01 | AVSNVIASLTCGRRF | 168.09 |
| HLA-DRB1*03:01 |  | 257.26 |
| HLA-DRB1*07:01 | AWGLLLMILHPDVQR | 389.63 |
| HLA-DRB1*03:01 |  | 688.11 |
| HLA-DRB1*07:01 | AWREQRRFSVSTLRN | 272.18 |
| HLA-DRB1*07:01 | AWTPVVVLNGLAAVR | 253.48 |
| HLA-DRB1*03:01 |  | 488.88 |
| HLA-DRB1*07:01 | CAAFANHSGRPFRPN | 242.34 |
| HLA-DRB1*03:01 | CGRRFEYDDPRFLRL | 170.30 |
| HLA-DRB1*07:01 |  | 682.85 |
| HLA-DRB1*07:01 | CLCAAFANHSGRPFR | 163.15 |
| HLA-DRB1*07:01 | DAQGHFVKPEAFLPF | 472.50 |
| HLA-DRB1*07:01 | DIEVQGFRIPKGTTL | 214.37 |
| HLA-DRB1*07:01 | DIVPLGVTHMTSRDI | 163.42 |
| HLA-DRB1*07:01 | DLFSAGMVTTSTTLA | 188.85 |
| HLA-DRB1*07:01 | DVFSLQLAWTPVVVL | 29.80 |
| HLA-DRB1*07:01 | EAACLCAAFANHSGR | 473.77 |
| HLA-DRB1*07:01 | EAFLPFSAGRRACLG | 168.92 |
| HLA-DRB1*07:01 | EALVPLAVIVAIFLL | 984.57 |
| HLA-DRB1*07:01 | EEAACLCAAFANHSG | 614.10 |
| HLA-DRB1*07:01 | EESGFLREVLNAVPV | 233.53 |
| HLA-DRB1*03:01 | EGLKEESGFLREVLN | 662.53 |
| HLA-DRB1*07:01 | ELFLFFTSLLQHFSF | 355.70 |
| HLA-DRB1*07:01 | EQRRFSVSTLRNLGL | 28.17 |
| HLA-DRB1*07:01 | ESGFLREVLNAVPVL | 189.95 |
| HLA-DRB1*07:01 |  | 165.27 |
| HLA-DRB1*07:01 | EVQGFRIPKGTTLIT | 112.72 |
| HLA-DRB1*07:01 | EVQRFGDIVPLGVTH | 209.23 |
| HLA-DRB1*07:01 | FAFLVSPSPYELCAV | 105.91 |
| HLA-DRB1*07:01 | FFTSLLQHFSFSVPT | 32.14 |
| HLA-DRB1*07:01 | FGDIVPLGVTHMTSR | 522.95 |
| HLA-DRB1*07:01 | FGDVFSLQLAWTPVV | 87.74 |
| HLA-DRB1*07:01 | FLFFTSLLQHFSFSV | 46.13 |
| HLA-DRB1*03:01 | FLLLVDLMHRRQRWA | 37.57 |
| HLA-DRB1*07:01 |  | 857.18 |
| HLA-DRB1*07:01 | FLPFSAGRRACLGEP | 437.15 |
| HLA-DRB1*07:01 |  | 59.73 |
| HLA-DRB1*03:01 | FLREVLNAVPVLLHI | 791.35 |
| HLA-DRB1*07:01 | FLVSPSPYELCAVPR | 824.36 |
| HLA-DRB1*07:01 | FQKAFLTQLDELLTE | 518.48 |
| HLA-DRB1*07:01 | FRIPKGTTLITNLSS | 713.96 |
| HLA-DRB1*07:01 | FSAGMVTTSTTLAWG | 163.35 |
| HLA-DRB1*07:01 | FSLQLAWTPVVVLNG | 24.15 |
| HLA-DRB1*07:01 | FSVSTLRNLGLGKKS | 337.67 |
| HLA-DRB1*07:01 | FTSLLQHFSFSVPTG | 34.95 |
| HLA-DRB1*07:01 | FVKPEAFLPFSAGRR | 279.60 |
| HLA-DRB1*07:01 | GDVFSLQLAWTPVVV | 68.54 |
| HLA-DRB1*07:01 | GFLREVLNAVPVLLH | 60.30 |
| HLA-DRB1*07:01 | GFRIPKGTTLITNLS | 324.84 |
| HLA-DRB1*07:01 | GHFVKPEAFLPFSAG | 451.36 |
| HLA-DRB1*07:01 | GLAAVREALVTHGED | 674.54 |
| HLA-DRB1*07:01 | GLEALVPLAVIVAIF | 408.82 |
| HLA-DRB1*03:01 | GLGNLLHVDFQNTPY | 607.67 |
| HLA-DRB1*03:01 | GLLLMILHPDVQRRV | 93.46 |
| HLA-DRB1*07:01 |  | 276.29 |
| HLA-DRB1*07:01 | GMVTTSTTLAWGLLL | 179.77 |
| HLA-DRB1*03:01 | GNLLHVDFQNTPYCF | 505.63 |
| HLA-DRB1*07:01 | GPAWREQRRFSVSTL | 533.67 |
| HLA-DRB1*03:01 | GRRFEYDDPRFLRLL | 131.87 |
| HLA-DRB1*07:01 |  | 512.68 |
| HLA-DRB1*07:01 | GTTLITNLSSVLKDE | 181.80 |
| HLA-DRB1*03:01 |  | 621.72 |
| HLA-DRB1*07:01 | GVTHMTSRDIEVQGF | 415.23 |
| HLA-DRB1*07:01 | HEVQRFGDIVPLGVT | 214.21 |
| HLA-DRB1*07:01 | HFVKPEAFLPFSAGR | 543.08 |
| HLA-DRB1*07:01 | HMTSRDIEVQGFRIP | 278.99 |
| HLA-DRB1*07:01 | IEVQGFRIPKGTTLI | 114.85 |
| HLA-DRB1*03:01 | IFLLLVDLMHRRQRW | 32.54 |
| HLA-DRB1*07:01 |  | 750.10 |
| HLA-DRB1*03:01 | ILHPDVQRRVQQEID | 297.28 |
| HLA-DRB1*07:01 | IPKGTTLITNLSSVL | 161.06 |
| HLA-DRB1*03:01 | ITNLSSVLKDEAVWE | 651.72 |
| HLA-DRB1*03:01 | IVAIFLLLVDLMHRR | 77.94 |
| HLA-DRB1*07:01 |  | 840.56 |
| HLA-DRB1*07:01 | IVPLGVTHMTSRDIE | 205.56 |
| HLA-DRB1*07:01 | KEESGFLREVLNAVP | 395.29 |
| HLA-DRB1*07:01 | KGTTLITNLSSVLKD | 130.81 |
| HLA-DRB1*03:01 |  | 569.84 |
| HLA-DRB1*07:01 | KPEAFLPFSAGRRAC | 373.03 |
| HLA-DRB1*03:01 | LAQEGLKEESGFLRE | 820.69 |
| HLA-DRB1*07:01 | LAWGLLLMILHPDVQ | 614.79 |
| HLA-DRB1*07:01 | LAWTPVVVLNGLAAV | 230.01 |
| HLA-DRB1*07:01 | LCAAFANHSGRPFRP | 200.71 |
| HLA-DRB1*03:01 | LDELLTEHRMTWDPA | 687.45 |
| HLA-DRB1*07:01 | LEALVPLAVIVAIFL | 653.30 |
| HLA-DRB1*07:01 | LEQWVTEEAACLCAA | 882.86 |
| HLA-DRB1*07:01 | LFFTSLLQHFSFSVP | 34.87 |
| HLA-DRB1*07:01 | LFLFFTSLLQHFSFS | 387.27 |
| HLA-DRB1*07:01 | LFSAGMVTTSTTLAW | 138.81 |
| HLA-DRB1*03:01 | LGNLLHVDFQNTPYC | 705.70 |
| HLA-DRB1*07:01 | LGVTHMTSRDIEVQG | 191.37 |
| HLA-DRB1*07:01 | LITNLSSVLKDEAVW | 595.33 |
| HLA-DRB1*07:01 | LKEESGFLREVLNAV | 565.99 |
| HLA-DRB1*03:01 | LLLMILHPDVQRRVQ | 62.49 |
| HLA-DRB1*07:01 |  | 364.35 |
| HLA-DRB1*03:01 | LLLVDLMHRRQRWAA | 56.79 |
| HLA-DRB1*03:01 |  | 59.81 |
| HLA-DRB1*07:01 | LLMILHPDVQRRVQQ | 740.79 |
| HLA-DRB1*07:01 | LLQHFSFSVPTGQPR | 181.10 |
| HLA-DRB1*03:01 | LLVDLMHRRQRWAAR | 191.73 |
| HLA-DRB1*03:01 | LMILHPDVQRRVQQE | 73.51 |
| HLA-DRB1*07:01 | LNAVPVLLHIPALAG | 129.93 |
| HLA-DRB1*07:01 | LNGLAAVREALVTHG | 248.79 |
| HLA-DRB1*07:01 | LPFSAGRRACLGEPL | 994.65 |
| HLA-DRB1*07:01 | LQHFSFSVPTGQPRP | 386.59 |
| HLA-DRB1*07:01 | LQLAWTPVVVLNGLA | 42.89 |
| HLA-DRB1*07:01 | LREVLNAVPVLLHIP | 59.41 |
| HLA-DRB1*03:01 |  | 758.36 |
| HLA-DRB1*07:01 | LRFQKAFLTQLDELL | 108.55 |
| HLA-DRB1*07:01 | LRRRFGDVFSLQLAW | 32.63 |
| HLA-DRB1*03:01 | LSSVLKDEAVWEKPF | 177.69 |
| HLA-DRB1*03:01 | LTCGRRFEYDDPRFL | 646.40 |
| HLA-DRB1*03:01 | LTQLDELLTEHRMTW | 817.09 |
| HLA-DRB1*07:01 | MELFLFFTSLLQHFS | 246.43 |
| HLA-DRB1*07:01 | MGLEALVPLAVIVAI | 333.87 |
| HLA-DRB1*03:01 | MILHPDVQRRVQQEI | 132.91 |
| HLA-DRB1*07:01 | MTSRDIEVQGFRIPK | 226.16 |
| HLA-DRB1*07:01 | MVTTSTTLAWGLLLM | 212.31 |
| HLA-DRB1*07:01 | NAVPVLLHIPALAGK | 109.95 |
| HLA-DRB1*07:01 | NGLAAVREALVTHGE | 460.37 |
| HLA-DRB1*03:01 | NLLHVDFQNTPYCFD | 845.84 |
| HLA-DRB1*03:01 | NLSSVLKDEAVWEKP | 247.87 |
| HLA-DRB1*03:01 | NPESSFNDENLRIVV | 867.30 |
| HLA-DRB1*07:01 | NVIASLTCGRRFEYD | 211.95 |
| HLA-DRB1*03:01 |  | 514.63 |
| HLA-DRB1*07:01 | PAWREQRRFSVSTLR | 269.00 |
| HLA-DRB1*07:01 | PEAFLPFSAGRRACL | 164.54 |
| HLA-DRB1*03:01 | PESSFNDENLRIVVA | 719.78 |
| HLA-DRB1*07:01 | PKGTTLITNLSSVLK | 102.33 |
| HLA-DRB1*03:01 |  | 596.01 |
| HLA-DRB1*07:01 | PLGVTHMTSRDIEVQ | 174.41 |
| HLA-DRB1*07:01 | PVPITQILGFGPRSQ | 835.27 |
| HLA-DRB1*07:01 | PVVVLNGLAAVREAL | 250.22 |
| HLA-DRB1*03:01 |  | 340.61 |
| HLA-DRB1*03:01 | QEGLKEESGFLREVL | 428.65 |
| HLA-DRB1*07:01 | QGFRIPKGTTLITNL | 189.55 |
| HLA-DRB1*07:01 | QGHFVKPEAFLPFSA | 340.42 |
| HLA-DRB1*07:01 | QHFSFSVPTGQPRPS | 851.25 |
| HLA-DRB1*07:01 | QLAWTPVVVLNGLAA | 83.06 |
| HLA-DRB1*03:01 | QLDELLTEHRMTWDP | 750.61 |
| HLA-DRB1*07:01 | QRFGDIVPLGVTHMT | 222.30 |
| HLA-DRB1*07:01 | QRRFSVSTLRNLGLG | 33.24 |
| HLA-DRB1*07:01 | RDIEVQGFRIPKGTT | 698.63 |
| HLA-DRB1*07:01 | REQRRFSVSTLRNLG | 32.64 |
| HLA-DRB1*07:01 | REVLNAVPVLLHIPA | 71.06 |
| HLA-DRB1*03:01 |  | 771.27 |
| HLA-DRB1*03:01 | RFEYDDPRFLRLLDL | 376.18 |
| HLA-DRB1*07:01 |  | 752.54 |
| HLA-DRB1*07:01 | RFGDIVPLGVTHMTS | 347.85 |
| HLA-DRB1*07:01 | RFGDVFSLQLAWTPV | 177.43 |
| HLA-DRB1*07:01 | RFQKAFLTQLDELLT | 198.62 |
| HLA-DRB1*07:01 | RFSVSTLRNLGLGKK | 96.08 |
| HLA-DRB1*07:01 | RIPKGTTLITNLSSV | 633.16 |
| HLA-DRB1*07:01 | RMELFLFFTSLLQHF | 252.90 |
| HLA-DRB1*07:01 | RPPVPITQILGFGPR | 952.81 |
| HLA-DRB1*03:01 | RRFEYDDPRFLRLLD | 170.15 |
| HLA-DRB1*07:01 |  | 665.78 |
| HLA-DRB1*07:01 | RRFGDVFSLQLAWTP | 67.99 |
| HLA-DRB1*07:01 | RRFSVSTLRNLGLGK | 46.33 |
| HLA-DRB1*07:01 | RRRFGDVFSLQLAWT | 33.23 |
| HLA-DRB1*07:01 | SAGMVTTSTTLAWGL | 147.37 |
| HLA-DRB1*07:01 | SGFLREVLNAVPVLL | 80.16 |
| HLA-DRB1*07:01 | SLLQHFSFSVPTGQP | 78.30 |
| HLA-DRB1*07:01 | SLQLAWTPVVVLNGL | 25.39 |
| HLA-DRB1*07:01 | SNVIASLTCGRRFEY | 139.75 |
| HLA-DRB1*03:01 |  | 347.57 |
| HLA-DRB1*07:01 | SRDIEVQGFRIPKGT | 403.67 |
| HLA-DRB1*03:01 | SSVLKDEAVWEKPFR | 297.02 |
| HLA-DRB1*07:01 | STTLAWGLLLMILHP | 659.28 |
| HLA-DRB1*03:01 | SVLKDEAVWEKPFRF | 510.42 |
| HLA-DRB1*07:01 | SVSTLRNLGLGKKSL | 885.27 |
| HLA-DRB1*03:01 | TCGRRFEYDDPRFLR | 276.15 |
| HLA-DRB1*07:01 | TEEAACLCAAFANHS | 729.38 |
| HLA-DRB1*07:01 | THMTSRDIEVQGFRI | 274.27 |
| HLA-DRB1*07:01 | TLAWGLLLMILHPDV | 642.97 |
| HLA-DRB1*07:01 | TLITNLSSVLKDEAV | 287.07 |
| HLA-DRB1*03:01 | TNLSSVLKDEAVWEK | 267.22 |
| HLA-DRB1*07:01 | TPVVVLNGLAAVREA | 254.53 |
| HLA-DRB1*03:01 |  | 311.20 |
| HLA-DRB1*03:01 | TQLDELLTEHRMTWD | 739.79 |
| HLA-DRB1*07:01 | TSLLQHFSFSVPTGQ | 45.76 |
| HLA-DRB1*07:01 | TSRDIEVQGFRIPKG | 281.08 |
| HLA-DRB1*07:01 | TSTTLAWGLLLMILH | 667.99 |
| HLA-DRB1*07:01 | TTLAWGLLLMILHPD | 849.95 |
| HLA-DRB1*07:01 | TTLITNLSSVLKDEA | 196.54 |
| HLA-DRB1*03:01 |  | 703.08 |
| HLA-DRB1*07:01 | TTSTTLAWGLLLMIL | 475.08 |
| HLA-DRB1*03:01 | VAIFLLLVDLMHRRQ | 45.89 |
| HLA-DRB1*07:01 |  | 773.03 |
| HLA-DRB1*07:01 | VFSLQLAWTPVVVLN | 23.00 |
| HLA-DRB1*07:01 | VIASLTCGRRFEYDD | 326.19 |
| HLA-DRB1*07:01 | VKPEAFLPFSAGRRA | 283.90 |
| HLA-DRB1*07:01 | VLNAVPVLLHIPALA | 162.30 |
| HLA-DRB1*07:01 | VLNGLAAVREALVTH | 185.02 |
| HLA-DRB1*07:01 | VPLGVTHMTSRDIEV | 155.11 |
| HLA-DRB1*07:01 | VQGFRIPKGTTLITN | 126.22 |
| HLA-DRB1*07:01 | VQRFGDIVPLGVTHM | 141.88 |
| HLA-DRB1*07:01 | VSNVIASLTCGRRFE | 161.52 |
| HLA-DRB1*03:01 |  | 254.32 |
| HLA-DRB1*07:01 | VTTSTTLAWGLLLMI | 243.62 |
| HLA-DRB1*07:01 | VVLNGLAAVREALVT | 162.33 |
| HLA-DRB1*07:01 | VVVLNGLAAVREALV | 144.61 |
| HLA-DRB1*03:01 |  | 512.77 |
| HLA-DRB1*07:01 | WGLLLMILHPDVQRR | 304.02 |
| HLA-DRB1*03:01 |  | 336.01 |
| HLA-DRB1*07:01 | WREQRRFSVSTLRNL | 30.71 |
| HLA-DRB1*07:01 |  | 288.28 |
| HLA-DRB1*03:01 | WTPVVVLNGLAAVRE | 393.05 |

**Supplementary table 3.** Shortlisted tolerogenic MHC II specific CD4+ T cell epitopes in CYP 2D6 autoantigen for vaccine design.

| Allele | Peptide Sequence | IC50 | Percentile Rank | IL-10 inducer | IFNg inducer | Allergenicity | Toxicity |
| --- | --- | --- | --- | --- | --- | --- | --- |
| H2-IAb | VGDLFSAGMVTTSTT | 227.03 | 0.86 | IL-10 inducer  (0.582) | Negative  (-0.66281506) | Non-allergen | Non-toxin |
| H2-IAb | LFSAGMVTTSTTLSW | 571.86 | 2.80 | IL-10 inducer  (0.512) | Negative  (-0.40982448) | Non-allergen | Non-toxin |

**Supplementary table 4.** Shortlisted tolerogenic HLA II specific CD4+ T cell epitopes in CYP 2D6 autoantigen for vaccine design.

| **Allele** | **Epitope** | **Region** | **IC50** | **Percentile Rank** | **IL-10**  **inducer** | **IFNg inducer** | **Allergenicity** | **Toxin** |
| --- | --- | --- | --- | --- | --- | --- | --- | --- |
| HLA-DRB1*07:01 | ADLFSAGMVTTSTTL | 300-314 | 314.93 | 24 | IL10 inducer  (0.522) | Negative  -0.13359248 | Non-Allergen | Non-toxin |
| HLA-DRB1*07:01 | AFLVSPSPYELCAVP | 482-496 | 242.63 | 19 | L10 inducer  (0.493) | Negative  -0.49429634 | Non-Allergen | Non-toxin |
| HLA-DRB1*07:01 | AGMVTTSTTLAWGLL | 305-319 | 176.51 | 15 | IL10 inducer  (0.455) | Negative  -0.07149604 | Non-Allergen | Non-toxin |
| HLA-DRB1*03:01 | AIFLLLVDLMHRRQR | 14-28 | 30.40 | 0.38 | IL10 inducer  (0.685) | Negative  -0.28895759 | Non-Allergen | Non-toxin |
| HLA-DRB1*07:01 |  |  | 713.47 | 39 | IL10 inducer  (0.685) | Negative  -0.28895759 | Non-Allergen | Non-toxin |
| HLA-DRB1*07:01 | AVPVLLHIPALAGKV | 226-240 | 57.96 | 4.90 | IL10 inducer  (0.575) | Negative  -0.025192334 | Non-Allergen | Non-toxin |
| HLA-DRB1*07:01 | AWREQRRFSVSTLRN | 127-141 | 272.18 | 21 | IL10 inducer  (0.51) | Negative  -0.17447177 | Non-Allergen | Non-toxin |
| HLA-DRB1*03:01 | CGRRFEYDDPRFLRL | 191-205 | 170.30 | 4 | IL10 inducer  (0.613) | Negative  -0.38729064 | Non-Allergen | Non-toxin |
| HLA-DRB1*07:01 |  |  | 682.85 | 38 | IL10 inducer  (0.612) | Negative  -0.38729064 | Non-Allergen | Non-toxin |
| HLA-DRB1*03:01 | DENLRIVVADLFSAG | 292-306 | 803.79 | 16 | IL10 inducer  (0.572) | Negative  -0.0069350174 | Non-Allergen | Non-toxin |
| HLA-DRB1*07:01 |  |  | 290.25 | 22 | IL10 inducer  (0.572) | Negative  -0.0069350174 | Non-Allergen | Non-toxin |
| HLA-DRB1*07:01 | DIEVQGFRIPKGTTL | 381-395 | 214.37 | 17 | L10 inducer  (0.388) | Negative  -0.46620572 | Non-Allergen | Non-toxin |
| HLA-DRB1*07:01 | ELFLFFTSLLQHFSF | 452-466 | 355.70 | 26 | IL10 inducer  (0.658) | Negative  -0.31977655 | Non-Allergen | Non-toxin |
| HLA-DRB1*07:01 | EVQRFGDIVPLGVTH | 362-376 | 209.23 | 17 | L10 inducer  (0.677) | Negative  -0.44527386 | Non-Allergen | Non-toxin |
| HLA-DRB1*07:01 | FAFLVSPSPYELCAV | 481-495 | 105.91 | 9.40 | IL10 inducer  (0.558) | Negative  -0.46786318 | Non-Allergen | Non-toxin |
| HLA-DRB1*07:01 | FFTSLLQHFSFSVPT | 456-470 | 32.14 | 2.30 | IL10 inducer  (0.66) | Negative  -0.54146309 | Non-Allergen | Non-toxin |
| HLA-DRB1*07:01 | FGDIVPLGVTHMTSR | 366-380 | 522.95 | 33 | IL10 inducer  (0.603) | Negative  -0.94014879 | Non-Allergen | Non-toxin |
| HLA-DRB1*07:01 |  |  | 87.74 | 7.90 | IL10 inducer  (0.598) | Negative  -0.2870051 | Non-Allergen | Non-toxin |
| HLA-DRB1*07:01 | FLFFTSLLQHFSFSV | 454-468 | 46.13 | 3.90 | IL10 inducer  (0.675) | Negative  -0.21930818 | Non-Allergen | Non-toxin |
| HLA-DRB1*07:01 | FLPFSAGRRACLGEP | 433-447 | 437.15 | 29 | IL10 inducer  (0.532) | Negative  -0.18451165 | Non-Allergen | Non-toxin |
| HLA-DRB1*07:01 | FLVSPSPYELCAVPR | 483-497 | 824.36 | 42 | IL10 inducer  (0.503) | Negative  -0.32204596 | Non-Allergen | Non-toxin |
| HLA-DRB1*07:01 | FRIPKGTTLITNLSS | 387-401 | 713.96 | 39 | IL10 inducer  (0.428) | Negative  -0.67722092 | Non-Allergen | Non-toxin |
| HLA-DRB1*07:01 | FSAGMVTTSTTLAWG | 303-317 | 163.35 | 14 | IL10 inducer  (0.487) | Negative  -0.45474223 | Non-Allergen | Non-toxin |
| HLA-DRB1*07:01 | FSLQLAWTPVVVLNG | 69-83 | 24.15 | 1.60 | IL10 inducer  (0.373) | Negative  -0.19601641 | Non-Allergen | Non-toxin |
| HLA-DRB1*07:01 | FTSLLQHFSFSVPTG | 457-471 | 34.95 | 2.60 | IL10 inducer  (0.64) | Negative  -0.44724792 | Non-Allergen | Non-toxin |
| HLA-DRB1*07:01 | GFRIPKGTTLITNLS | 386-400 | 324.84 | 24 | IL10 inducer  (0.427) | Negative  -0.65701274 | Non-Allergen | Non-toxin |
| HLA-DRB1*07:01 | GHFVKPEAFLPFSAG | 425-439 | 451.36 | 30 | IL10 inducer  (0.493) | Negative  -0.049324136 | Non-Allergen | Non-toxin |
| HLA-DRB1*07:01 | GPRSQGVFLARYGPA | 113-127 | 987.40 | 47 | IL10 inducer  (0.325) | Negative  -0.016483882 | Non-Allergen | Non-toxin |
| HLA-DRB1*03:01 | GRRFEYDDPRFLRLL | 192-206 | 131.87 | 3.20 | IL10 inducer  (0.628) | Negative  -0.18459203 | Non-Allergen | Non-toxin |
| HLA-DRB1*07:01 |  |  | 512.68 | 32 | IL10 inducer  (0.628) | Negative  -0.18459203 | Non-Allergen | Non-toxin |
| HLA-DRB1*07:01 | HEVQRFGDIVPLGVT | 361-375 | 214.21 | 17 | L10 inducer  (0.648) | Negative  -0.19236111 | Non-Allergen | Non-toxin |
| HLA-DRB1*07:01 | HMPYTTAVIHEVQRF | 352-366 | 133.34 | 12 | IL10 inducer  (0.682) | Negative  -0.37817695 | Non-Allergen | Non-toxin |
| HLA-DRB1*07:01 | HMTSRDIEVQGFRIP | 376-390 | 278.99 | 21 | IL10 inducer  (0.477) | Negative  -1.0919645 | Non-Allergen | Non-toxin |
| HLA-DRB1*03:01 | IFLLLVDLMHRRQRW | 15-29 | 32.54 | 0.44 | IL10 inducer  (0.687) | Negative  -0.051581412 | Non-Allergen | Non-toxin |
| HLA-DRB1*07:01 |  |  | 750.10 | 40 | IL10 inducer  (0.687) | Negative  -0.051581412 | Non-Allergen | Non-toxin |
| HLA-DRB1*03:01 | IVAIFLLLVDLMHRR | 12-26 | 77.94 | 2 | IL10 inducer  (0.635) | Negative  -0.21911829 | Non-Allergen | Non-toxin |
| HLA-DRB1*07:01 |  |  | 840.56 | 43 | IL10 inducer  (0.655) | Negative  -0.21911829 | Non-Allergen | Non-toxin |
| HLA-DRB1*07:01 | IVPLGVTHMTSRDIE | 369-383 | 205.56 | 17 | IL10 inducer  (0.592) | Negative  -0.95158969 | Non-Allergen | Non-toxin |
| HLA-DRB1*07:01 | IVVADLFSAGMVTTS | 297-311 | 364.79 | 26 | IL10 inducer  (0.523) | Negative  -0.62295981 | Non-Allergen | Non-toxin |
| HLA-DRB1*03:01 | KGTTLITNLSSVLKD | 391-405 | 569.84 | 12 | IL10 inducer  (0.51) | Negative  -0.52545771 | Non-Allergen | Non-toxin |
| HLA-DRB1*07:01 |  |  | 130.81 | 12 | IL10 inducer  (0.51) | Negative  -0.52545771 | Non-Allergen | Non-toxin |
| HLA-DRB1*07:01 | KVLRFQKAFLTQLDE | 239-253 | 37.48 | 2.90 | IL10 inducer  (0.693) | Negative  -0.36600417 | Non-Allergen | Non-toxin |
| HLA-DRB1*07:01 | LFFTSLLQHFSFSVP | 455-469 | 34.87 | 2.60 | IL10 inducer  (0.612) | Negative  -0.56155191 | Non-Allergen | Non-toxin |
| HLA-DRB1*07:01 | LFLFFTSLLQHFSFS | 453-467 | 387.27 | 27 | IL10 inducer  (0.627) | Negative  -0.040108823 | Non-Allergen | Non-toxin |
| HLA-DRB1*07:01 | LFSAGMVTTSTTLAW | 302-316 | 138.81 | 12 | IL10 inducer  (0.488) | Negative  -0.2727525 | Non-Allergen | Non-toxin |
| HLA-DRB1*03:01 | LGNLLHVDFQNTPYC | 43-57 | 705.70 | 14 | IL10 inducer  (0.572) | Negative  -0.0155045 | Non-Allergen | Non-toxin |
| HLA-DRB1*03:01 | LHIPALAGKVLRFQK | 231-245 | 705.32 | 14 | IL10 inducer  (0.647) | Negative  -0.21482126 | Non-Allergen | Non-toxin |
| HLA-DRB1*07:01 |  |  | 183.59 | 15 | IL10 inducer  (0.647 | Negative  -0.21482126 | Non-Allergen | Non-toxin |
| HLA-DRB1*07:01 | LITNLSSVLKDEAVW | 395-409 | 595.33 | 35 | IL10 inducer  (0.368) | Negative  -0.4019765 | Non-Allergen | Non-toxin |
| HLA-DRB1*03:01 | LLHIPALAGKVLRFQ | 230-244 | 928.88 | 18 | IL10 inducer  (0.623) | Negative  -0.068026415 | Non-Allergen | Non-toxin |
| HLA-DRB1*07:01 |  |  | 100.08 | 9 | IL10 inducer  (0.623) | Negative  -0.068026415 | Non-Allergen | Non-toxin |
| HLA-DRB1*07:01 | LLQHFSFSVPTGQPR | 460-474 | 181.10 | 15 | IL10 inducer  (0.615) | Negative  -0.10802673 | Non-Allergen | Non-toxin |
| HLA-DRB1*07:01 | LNAVPVLLHIPALAG | 224-238 | 129.93 | 12 | IL10 inducer  (0.457) | Negative  -0.36440417 | Non-Allergen | Non-toxin |
| HLA-DRB1*07:01 | LQHFSFSVPTGQPRP | 461-475 | 386.59 | 27 | IL10 inducer  (0.607) | Negative  -0.16337351 | Non-Allergen | Non-toxin |
| HLA-DRB1*07:01 | LRFQKAFLTQLDELL | 241-255 | 108.55 | 9.60 | IL10 inducer  (0.69) | Negative  -0.025403035 | Non-Allergen | Non-toxin |
| HLA-DRB1*03:01 | LRIVVADLFSAGMVT | 295-309 | 755.01 | 15 | IL10 inducer  (0.538) | Negative  -0.30525199 | Non-Allergen | Non-toxin |
| HLA-DRB1*07:01 |  |  | 389.48 | 27 | IL10 inducer  (0.548) | Negative  -0.30525199 | Non-Allergen | Non-toxin |
| HLA-DRB1*03:01 | LTCGRRFEYDDPRFL | 189-203 | 646.40 | 13 | IL10 inducer  (0.623) | Negative  -0.24096184 | Non-Allergen | Non-toxin |
| HLA-DRB1*07:01 | MGDQAHMPYTTAVIH | 347-361 | 873.11 | 44 | IL10 inducer  (0.653) | Negative  -0.29828068 | Non-Allergen | Non-toxin |
| HLA-DRB1*07:01 | MTSRDIEVQGFRIPK | 377-391 | 226.16 | 18 | L10 inducer  (0.49) | Negative  -0.7686883 | Non-Allergen | Non-toxin |
| HLA-DRB1*07:01 | NAVPVLLHIPALAGK | 225-239 | 109.95 | 9.70 | (IL10 inducer  0.542) | Negative  -0.17367147 | Non-Allergen | Non-toxin |
| HLA-DRB1*03:01 | NLLHVDFQNTPYCFD | 45-59 | 845.84 | 17 | IL10 inducer  0.643 | Negative  -0.38583963 | Non-Allergen | Non-toxin |
| HLA-DRB1*03:01 | NLRIVVADLFSAGMV | 294-308 | 708.28 | 14 | IL10 inducer  (0.528) | Negative  -0.11686082 | Non-Allergen | Non-toxin |
| HLA-DRB1*07:01 |  |  | 277.62 | 21 | IL10 inducer  (0.528) | Negative  -0.11686082 | Non-Allergen | Non-toxin |
| HLA-DRB1*03:01 | NLSSVLKDEAVWEKP | 398-412 | 247.87 | 5.60 | L10 inducer  (0.418) | Negative  -0.29621597 | Non-Allergen | Non-toxin |
| HLA-DRB1*07:01 | PLGVTHMTSRDIEVQ | 371-385 | 174.41 | 15 | IL10 inducer  (0.593) | Negative  -0.57878935 | Non-Allergen | Non-toxin |
| HLA-DRB1*07:01 | PVPITQILGFGPRSQ | 103-117 | 835.27 | 43 | IL10 inducer  (0.478) | Negative  -0.21248555 | Non-Allergen | Non-toxin |
| HLA-DRB1*03:01 | QEGLKEESGFLREVL | 210-224 | 428.65 | 9 | IL10 inducer  0.543 | Negative  -0.27411742 | Non-Allergen | Non-toxin |
| HLA-DRB1*07:01 | QRFGDIVPLGVTHMT | 364-378 | 222.30 | 18 | L10 inducer  (0.635) | Negative  -0.70893138 | Non-Allergen | Non-toxin |
| HLA-DRB1*07:01 | RDIEVQGFRIPKGTT | 380-394 | 698.63 | 39 | IL10 inducer  (0.45) | Negative  -0.86266453 | Non-Allergen | Non-toxin |
| HLA-DRB1*07:01 | REQRRFSVSTLRNLG | 129-143 | 32.64 | 2.40 | IL10 inducer  (0.555) | Negative  -0.15733669 | Non-Allergen | Non-toxin |
| HLA-DRB1*03:01 | RFEYDDPRFLRLLDL | 194-208 | 376.18 | 7.90 | IL10 inducer  (0.652) | Negative  -0.33137368 | Non-Allergen | Non-toxin |
| HLA-DRB1*07:01 |  |  | 752.54 | 40 | IL10 inducer  (0.652) | Negative  -0.33137368 | Non-Allergen | Non-toxin |
| HLA-DRB1*07:01 | RFGDIVPLGVTHMTS | 365-379 | 347.85 | 25 | IL10 inducer  (0.625) | Negative  -1.1077686 | Non-Allergen | Non-toxin |
| HLA-DRB1*07:01 | RFGDVFSLQLAWTPV | 64-78 | 177.43 | 15 | IL10 inducer  (0.648) | Negative  -0.45504666 | Non-Allergen | Non-toxin |
| HLA-DRB1*03:01 | RIVVADLFSAGMVTT | 296-310 | 919.55 | 18 | IL10 inducer  (0.535) | Negative  -0.2302519 | Non-Allergen | Non-toxin |
| HLA-DRB1*07:01 |  |  | 397.04 | 28 | IL10 inducer  (0.535) | Negative  -0.2302519 | Non-Allergen | Non-toxin |
| HLA-DRB1*03:01 | RRFEYDDPRFLRLLD | 193-207 | 170.15 | 4 | IL10 inducer  (0.647) | Negative  -0.003691101 | Non-Allergen | Non-toxin |
| HLA-DRB1*07:01 |  |  | 665.78 | 38 | IL10 inducer  (0.652) | Negative  -0.003691101 | Non-Allergen | Non-toxin |
| HLA-DRB1*07:01 | SAGMVTTSTTLAWGL | 304-318 | 147.37 | 13 | IL10 inducer  (0.453) | Negative  -0.0020614912 | Non-Allergen | Non-toxin |
| HLA-DRB1*07:01 | SRDIEVQGFRIPKGT | 379-393 | 403.67 | 28 | IL10 inducer  (0.538) | Negative  -0.70288511 | Non-Allergen | Non-toxin |
| HLA-DRB1*03:01 | TCGRRFEYDDPRFLR | 190-204 | 276.15 | 6.10 | IL10 inducer  (0.623) | Negative  -0.1047457 | Non-Allergen | Non-toxin |
| HLA-DRB1*07:01 | THMTSRDIEVQGFRI | 375-389 | 274.27 | 21 | IL10 inducer  (0.535) | Negative  -0.75794627 | Non-Allergen | Non-toxin |
| HLA-DRB1*03:01 | TNLSSVLKDEAVWEK | 397-411 | 267.22 | 5.90 | IL10 inducer  (0.437) | Negative  -0.14124887 | Non-Allergen | Non-toxin |
| HLA-DRB1*07:01 | TSLLQHFSFSVPTGQ | 458-472 | 45.76 | 3.80 | IL10 inducer  (0.622) | Negative  -0.58536503 | Non-Allergen | Non-toxin |
| HLA-DRB1*07:01 | TSRDIEVQGFRIPKG | 378-392 | 281.08 | 22 | IL10 inducer  (0.493) | Negative  -0.7467795 | Non-Allergen | Non-toxin |
| HLA-DRB1*03:01 | TTLITNLSSVLKDEA | 393-407 | 703.08 | 14 | IL10 inducer  (0.428) | Negative  -0.5102756 | Non-Allergen | Non-toxin |
| HLA-DRB1*07:01 |  |  | 196.54 | 16 | IL10 inducer  (0.428) | Negative  -0.5102756 | Non-Allergen | Non-toxin |
| HLA-DRB1*07:01 | VADLFSAGMVTTSTT | 299-313 | 554.78 | 34 | IL10 inducer  (0.54) | Negative  -0.23123227 | Non-Allergen | Non-toxin |
| HLA-DRB1*03:01 | VAIFLLLVDLMHRRQ | 13-27 | 45.89 | 0.82 | IL10 inducer  (0.68) | Negative  -0.29809723 | Non-Allergen | Non-toxin |
| HLA-DRB1*07:01 |  |  | 773.03 | 41 | IL10 inducer  (0.68) | Negative  -0.29809723 | Non-Allergen | Non-toxin |
| HLA-DRB1*07:01 | VFSLQLAWTPVVVLN | 68-82 | 23.00 | 1.50 | IL10 inducer  (0.438) | Negative  -0.12803696 | Non-Allergen | Non-toxin |
| HLA-DRB1*07:01 | VLLHIPALAGKVLRF | 299-243 | 54.87 | 4.70 | IL10 inducer  (0.585) | Negative  -0.22284623 | Non-Allergen | Non-toxin |
| HLA-DRB1*07:01 | VLNAVPVLLHIPALA | 223-237 | 162.30 | 14 | IL10 inducer  (0.418) | Negative  -0.15738463 | Non-Allergen | Non-toxin |
| HLA-DRB1*07:01 | VLRFQKAFLTQLDEL | 240-254 | 57.95 | 4.90 | IL10 inducer  (0.683) | Negative  -0.25527112 | Non-Allergen | Non-toxin |
| HLA-DRB1*07:01 | VPLGVTHMTSRDIEV | 370-384 | 155.11 | 14 | IL10 inducer  (0.612) | Negative  -0.45114239 | Non-Allergen | Non-toxin |
| HLA-DRB1*07:01 | VQRFGDIVPLGVTHM | 363-377 | 141.88 | 13 | IL10 inducer  (0.625) | Negative  -0.58219957 | Non-Allergen | Non-toxin |
| HLA-DRB1*07:01 | VVADLFSAGMVTTST | 298-312 | 417.43 | 29 | IL10 inducer  (0.532) | Negative  -0.40044594 | Non-Allergen | Non-toxin |
| HLA-DRB1*07:01 | WREQRRFSVSTLRNL | 128-142 | 30.71 | 2.30 | IL10 inducer  (0.543) | Negative  -0.39382573 | Non-Allergen | Non-toxin |
